# DREAM-Yara: An exact read mapper for very large databases with short update time

**DOI:** 10.1101/256354

**Authors:** Temesgen Hailemariam Dadi, Enrico Siragusa, Vitor C. Piro, Andreas Andrusch, Enrico Seiler, Bernhard Y. Renard, Knut Reinert

## Abstract

**Motivation:** Mapping-based approaches have become limited in their application to very large sets of references since computing an FM-index for very large databases (e.g. *>* 10 GB) has become a bottleneck. This affects many analyses that need such index as an essential step for approximate matching of the NGS reads to reference databases. For instance, in typical metagenomics analysis, the size of the reference sequences has become prohibitive to compute a single full-text index on standard machines. Even on large memory machines, computing such index takes about one day of computing time. As a result, updates of indices are rarely performed. Hence, it is desirable to create an alternative way of indexing while preserving fast search times.

**Results:** To solve the index construction and update problem we propose the DREAM (Dynamic seaRchablE pArallel coMpressed index) framework and provide an implementation. The main contributions are the introduction of an approximate search distributor directories via a novel use of Bloom filters. We combine several Bloom filters to form an *interleaved* Bloom filter and use this new data structure to quickly exclude reads for parts of the databases where they cannot match. This allows us to keep the databases in several indices which can be easily rebuilt if parts are updated while maintaining a fast search time. The second main contribution is an implementation of DREAM-Yara a distributed version of a fully sensitive read mapper under the DREAM framework.

**Availability:** https://gitlab.com/pirovc/dream_yara/

## 1 Introduction

Within the last ten years, modern sequencing technologies have brought a super-exponential growth of sequencing capacities. This has enabled the cheap sequencing of the genomic content of pangenomes (Consortium, 2018), metagenomes, or many individuals of the same species (e.g. the 100,000 genome project) that differ only slightly from each other. Yet, the small individual differences are of interest (i.e. SNPs, or small structural polymorphisms) to elucidate the cause of diseases or reconstruct evolutionary events.

These datasets expose interesting characteristics. They are large, while some large fractions are highly redundant (e.g. 100,000 genome project, or storing different strains of bacteria) and hence amenable to compression techniques (e.g. Rahn *et al.* (2014); Schneeberger *et al.* (2009)). On the other hand, compression usually makes it costly to implement the main operations on the data, namely *finding approximate matches* of (many) queries (approximate in the sense of edit distance).

While a lot of research has focused on indexing such datasets, the resulting solutions lack, in general, the ability to easily *change* the underlying datasets. That means it is costly (a) to change small parts of a sequence, or (b) to add or delete complete sequences while maintaining the ability to support fast approximate string searches. For example, in metagenomics, this problem becomes more and more recurrent. Many metagenomics search tools (e.g. Hauswedell *et al.* (2014); Piro *et al.* (2016)) and read mappers Li and Durbin (2010); Siragusa (2013) use Burrows-Wheeler-Transform (BWT) based FM-indices (also often referred to as compressed suffix arrays (CSA) Ferragina and Manzini (2000)) which have to index about 50 to 200 gigabases. Due to constant database updates changes occur on a daily or weekly basis and thus require a newly constructed index. Recomputing a single index of this size is quite costly in terms of space and time, even if approaches of merging BWTs are used Bauer *et al.* (2011); Sirén (2009). For example, it takes about one day to compute the index for the dataset used in our experiments. On the other hand, the ability for fast approximate searches in such an index is crucial. It is used either directly to find all approximate occurrences of a (short) string or parts of it in seed-and-extend approaches.

In this work, we address the problem posed by relying on one large index. We propose a framework which can offer *various* solutions for the above areas depending on some key parameters of the input set (size of the input, amount of redundancy, the importance of rebuilding time vs. search time). We name the framework a DREAM index (Dynamic seaRchablE pArallel coMpressed index) and describe in this work a first working implementation of it which allows to easily update the underlying database and at the same time fast read mapping using a standard approach.

In a similar work, Mohamadi *et al.* (2015) presented a method where references are partitioned and indexed separately based on size. Then they used on-the-fly constructed Bloom filters to dispatch and map reads against the individual partitions. This approach comes short when reference datasets are rather large and there is a need to create many partitions. While investigating this method we noticed two major bottlenecks. First, read dispatching was inefficient, meaning almost all reads were being mapped against all partitions resulting in slower overall mapping time. In addition, the dispatched/partitioned reads are written to disc creating an additional IO overhead. Second, the merging step to create one alignment result from individual alignment files takes too long.

In the following, we describe the general DREAM index framework, followed by a description of our implementation which consists of three major contributions. First, a taxonomy based clustering/binning method for a collection of database sequences (e.g. bacterial genomes), second a novel data structure for quickly distributing reads to bins for mapping that relies on a combination of Bloom filters (Bloom, 1970) and *k*-mer counting, and lastly a distributed, parallel version of the Yara read mapper (Siragusa, 2013).

In spite of being published as a Ph.D. thesis in (Siragusa, 2013), we have also described and evaluated the standard Yara read mapper in detail. Yara is an exact read mapping tool that is efficient, easy to use, and produces well-defined and interpretable results. It outperforms previous exact tools like RazerS 3 (Weese *et al.*, 2012) by a factor of 200 and even heuristic methods like Bowtie 2 (Langmead and Salzberg, 2012) and BWA-backtrack (Li and Durbin, 2010) by two and three times. The efficiency of the tool is due to a novel combination of known algorithmic concepts and a solid implementation based on the SeqAn library (Döring *et al.*, 2008).

## 2 Methods

### 2.1 The DREAM index framework

In Figure 1 we describe the general DREAM index framework. On the left, we show the input to our framework, namely a set of database sequences that need to be indexed and one or more sets of sequencing reads. As pointed out above, we want to be able to add sequences to and (possibly) delete sequences from the index quickly. At the same time, we want to be able to conduct approximate searches of reads in the index. This is achieved by splitting the database sequences into smaller clusters which contain relatively similar sequences. Then each cluster is indexed with a method of choice, appropriate for the respective approximate search method. We address the resulting indices as sub-indices.

**Fig. 1.**
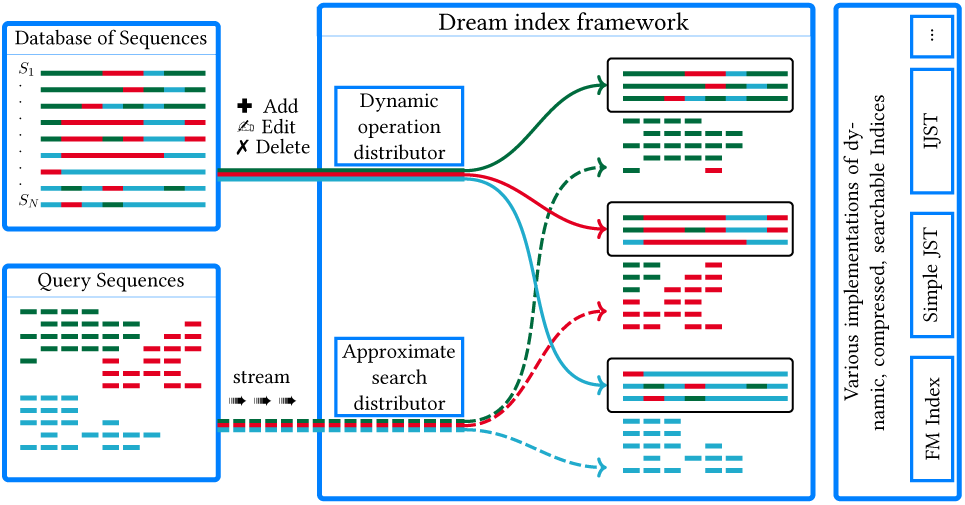
Sketch of the DREAM index framework. The red sequence piece among the green ones symbolizes that we do not require a perfect partitioning allowing us to use fast methods. The boxes on the right symbolize the potential use of different index implementations. Note that we use in this work solely FM-indices.

In more detail, the DREAM index consists of two layers for access and a collection of smaller sub-indices that contain highly similar (sub)sequences. The *dynamic operation distributor* layer will determine which sub-indices need to be created or changed to complete the update operations and trigger the respective operations in parallel on the sub-indices. The *approximate search distributor* layer will determine which sub-indices need to be searched, conduct the searches in parallel, and consolidate the results. It is obvious, that this will work better if we place relatively similar sequences (e.g. bacterial genomes of the same genus) into the same sub-index. Note that this will also benefit compression of those indices. However, we will not address compression deeper within this work.

We will rather use standard FM-indices for the sub-indices since they support fast approximate queries and give a coarse-grained dynamization by simply rebuilding a sub-index if needed. For other implementations of a sub-index, different solutions for the approximate search are possible and not within the scope of this publication.

For the dynamic operation and approximate search layer, we applied the *k*-mer counting lemma together with *k*-mer dictionaries based on a novel type of interleaved Bloom filters. Both strategies are discussed in the following sections.

### 2.2 Binning sequences, dynamic updates

For binning references, we used TaxSBP (https://github.com/pirovc/taxsbp), an implementation of the approximation algorithm for the hierarchically structured bin packing problem (Codenotti *et al.*, 2004) based on the NCBI Taxonomy database (Federhen, 2012). This clustering method is very efficient, given that it uses the "pre-clustered" taxonomic tree information to generate similarly sized groups of closely related sequences. In this work, we will consider contiguous sequences in the given reference genomes as the smallest unit of sequences that can be clustered into bins. That means we will not split those sequences into smaller parts. The results we present later are based on a metagenomics datasets for which it is relatively easy to obtain a taxonomic tree. Adding and removing sequences is also straightforward once their taxonomic classification is known. Note that in the absence of taxonomic information other, e.g. *k*-mer based, clustering methods can be used in our framework.

We assume now that we have divided the database text *T* into *b* bins in such a way, that a bin *B_i_* contains similar parts of *T*. For our approximate search distributor, we use what we call a *binning dictionary* in conjunction with a well-know *k*-mer counting lemma. The general idea of a binning dictionary *D* is that we will mark for a fixed *k*-mer in which bin it occurs using a *binning* bitvector, i.e we set the *i*-th bit to 1 if the *k*-mer is present in bin *B_i_*. Then, when we want to search a pattern *p* approximately, the following well-known Lemma gives a necessary condition for the pattern to occur in a bin.

Lemma 1. *For a given k and number of errors e, there are k_p_* =*| p | −k* + 1 *many k-mers in p and an approximate occurrence of p in T has to share at least t* = (*k_p_* − *k* • *e*) *k-mers*.

We now discuss how to compute a binning bitvector efficiently using a *global* data structure as opposed to the approach of Mohamadi *et al.* (2015) et al. who used *b* many Bloom filters.

### 2.3 Interleaved Bloom Filter

A Bloom filter is simply a bitvector of size *n* and a set of *h* independent hash functions that map a key value, in our case a *k*-mer, to one of the bit positions. To insert a *k*-mer to a Bloom filter we simply set *h* bit positions defined by *h* hash functions to 1. Collisions are allowed in the expense of false positive results. During lookup, a *k*-mer is considered present in the Bloom filter, if all *h* positions return a 1. Note that a Bloom filter can give a false positive answer if the *h* bits were set by other *k*-mers. However, if the Bloom filter size is large enough, the probability of a false positive answer is low. A Bloom filter of size *n* bits with *h* different hash functions and *m* elements inserted has the following probability of giving a false positive answer.

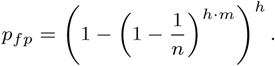

For this reason, we have to allocate sufficient space, such that *p_fp_* does not become too large.

One problem of using conventional Bloom filters is, that one cannot store directly a binning bitvector. A Bloom filter only answers set membership queries. Hence, Mohamadi *et al.* (2015) used a Bloom filter for each bin as opposed to resolving all bins simultaneously. To alleviate the problem, we propose to combine *b* Bloom filters (one for each bin) with *identical* hash functions into a *single* Bloom filter by interleaving them. To put it differently, we replace each bit in the single Bloom filter by a (sub)-bitvector of size *b*, where the *i*-th bit “belongs” to the Bloom filter for bin *B_i_*. We call the resulting Bloom filter an *Interleaved* Bloom Filter (IBF). The IBF has a size of *b · n*. When inserting a *k*-mer from bin *B_i_* into the IBF, we compute all *h* hash functions which point us to the position of the block where the sub-bitvectors are and then simply set the *i*-th bit from the respective beginnings. Hence, we effectively interleave *b* Bloom filters in a way that allows us to retrieve the binning bitvectors for the *h* hash functions easily. When querying in which bins a *k*-mer is, we would retrieve the *h* sub-bitvectors and apply a logical AND to them which results in the required binning bitvector indicating the membership of the *k*-mer in the bins. This approach has a significant advantage in query time as retrieving a (sub)-bitvector is extremely cache-friendly. The procedure is depicted in Figure 2.

**Fig. 2.**
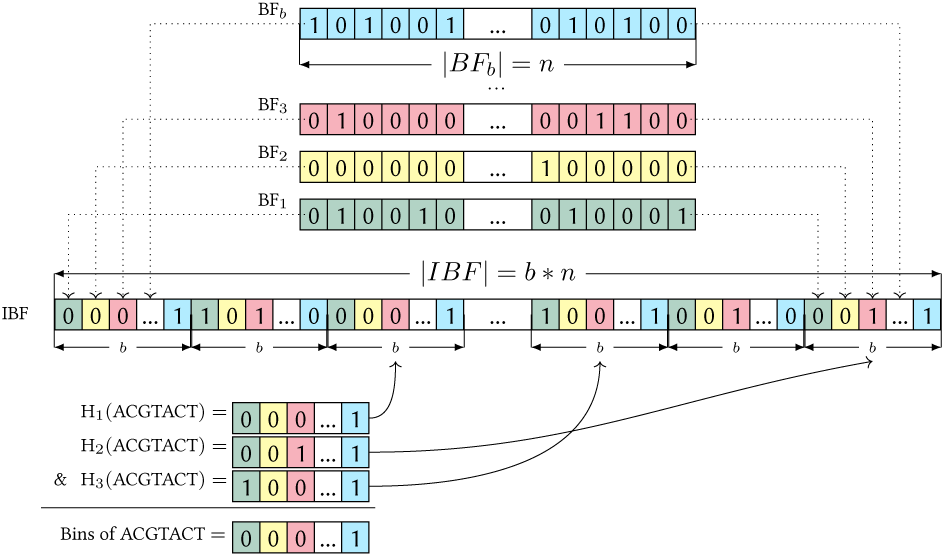
Example of an IBF. Differently colored Bloom filters of length *n* for the *b* bins are shown in the top. The individual Bloom filters are interleaved to make an IBF of size *b × n*. In the example, we retrieve 3 positions for a *k*-mer (ACGTACT) using 3 different hash functions. The corresponding sub bitvectors are combined with a bitwise & giving us the needed binning bitvector.

We anticipate that the IBF will be a very useful data structure for set membership of objects in bins and will see a wide usage, especially for assessing *k*-mer content. When writing this paper, it was brought to our attention that Bradley et al. independently thought of a similar data structure in Bradley *et al.* (2017), although they do not use them in conjunction with the *k*-mer Lemma and do not interleave them.

To decide which bins are a potential target for a read we apply Lemma 1 (the *k*-mer counting Lemma.) The IBF tells in which bin a given *k*-mer occurs by returning a binning bitvector. Hence, we can simply look up each *k*-mer from a pattern in the *IBF*, retrieve the binning bitvector marking its occurrences in the bins, and update an array of counters for each bin. If the counter exceeds the threshold for the bin, the pattern will be searched approximately in the bin, otherwise not. This approach is depicted in figure 3.

**Fig. 3.**
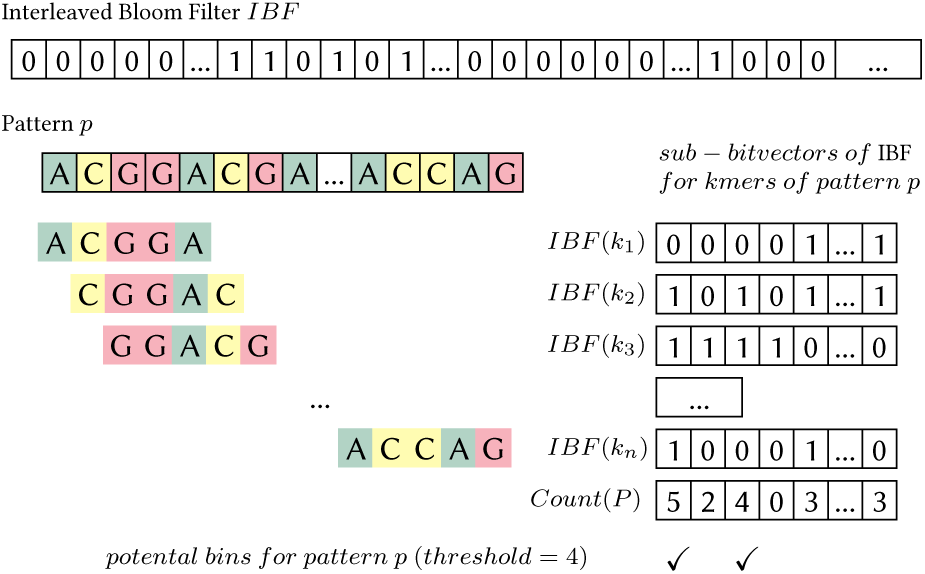
The k-mer counting Lemma using Interleaved Bloom Filter (IBF). For each *k*-mer *k_i_* generated from a pattern *p* we extract binning sub-bitvectors *SV*(*k_i_*) representing the bins containing k-mer ki. For all set bits in *SV*(*k_i_*) we increment the counter of corresponding bin. Bins whose counter is greater than or equals to the threshold (in this case 4) need to be validated for *p*.

The IBF can be partly updated in a straightforward way to reflect changes in bins. Consider the contents of the i^th^ bin has changed. First we reset the corresponding i^th^ bits from every sub-bitvector of the IBF. Then we add the kmers from the same updated bin to the IBF. This can be done in parallel for multiple affected bins.

### 2.4 The Yara read mapper

In this section, we describe Yara (Siragusa, 2013), currently the state-of-the art read mapper of the SeqAn library, an exact read mapping tool that is efficient, easy to use and produces well-defined and interpretable results. It outperforms previous exact tools like RazerS3 (Weese *et al.*, 2012) by a factor of 200 in speed and even heuristic methods like Bowtie 2 (Langmead and Salzberg, 2012) and BWA (Li and Durbin, 2010) by a factor of two and three, respectively (Siragusa, 2013). The efficiency of Yara is due to a novel combination of known algorithmic concepts and a solid implementation based on the SeqAn library (Reinert *et al.*, 2017).

Yara is based on the concept of best+*x* mapping which we advocate as more practical than conventional all-mapping (i.e. reporting all locations within a certain error bound). Mapping strata have been implemented also in Bowtie (Langmead *et al.*, 2009), RazerS3 (Weese *et al.*, 2012) and GEM (Marco-Sola *et al.*, 2012).

To explain, we define the *e*-stratum

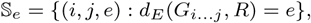

as the set of all mapping locations of a read *R* at edit distance *e* from the reference genome *G*. Conventional all-mapping reports all mapping locations within an absolute error threshold *k*

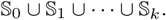

Best+*x* mapping restricts mapping locations to a fixed distance *x* from the *optimal* stratum *b*. That means if the distance of any optimal mapping location for read *R* is

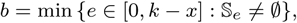

then best+*x* mapping considers only mapping locations within a *relative* suboptimality error threshold *x*

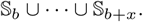

For example, suppose a read best-maps somewhere at edit distance *b* = 2 on the reference genome. Yara in mode best+0 reports all locations at distance 2, while in mode best+1 reports all locations at distance 2 and 3. Conversely, an all-mapper with absolute threshold *k* = 5 would report all locations up to distance 5.

To efficiently map reads within best+*x* strata, Yara combines three algorithmic ideas: adaptive filtration using approximate seeds, iterative pigeonhole filtration, greedy verification of candidate mapping locations.

Yara splits each read in *k* + 1 non-overlapping exact seeds and counts the number of candidate locations of each seed in the reference genome using an FM-index. According to the pigeonhole principle for approximate string matching, an all-mapper would verify all candidate locations of the *k* + 1 seeds and find all mapping locations at edit distance 0 to *k*. Yara instead proceeds seed by seed and determines *b* by verifying *b* + 1 seeds and checking that S_*b*_ is the first non-empty stratum. Iterating the pigeonhole principle, the tool stops after *b* + *x* + 1 seeds. As the order in which seeds are processed does not affect the results, Yara tries to minimize the number of verifications by processing the seeds greedily from the least to the most frequent seed. Moreover, Yara decides, based on fine-tuned internal thresholds, if it is worth proceeding with the verification phase or alternatively applying a stronger filtration scheme using 1 or 2-approximate seeds. The verification is done using a banded version of Myers bit-vector algorithm.

Yara maps paired-end reads independently (as single ends). After mapping, the tool selects a primary mapping location by pairing ends according to the estimated fragment size. Nonetheless, Yara reports all relevant mapping locations per end, as the lack of proper pairing signals potential structural variations.

Yara is already in use by several groups, it was used for improved metagenomics classification (Dadi *et al.*, 2017) and was tested favorably by (Břinda *et al.*, 2016). In addition it scales well using many threads. For those reasons we choose it as the first implementation within the DREAM framework and call the resulting tool DREAM-Yara.

### 2.5 Distributed-Yara

A straightforward approach to do a distributed mapping is to split the reference genome into smaller partitions, index them separately and search sequencing reads against each index. Such approach improve indexing times, but it creates an overhead on the mapping step. To make matters even worse the mapping results have to be consolidated, which is a complicated process as we need to consider the ranking of multiples mappings per read such as identifying primary mapping locations. Nevertheless, we applied this strategy using the Yara read mapper and named it *Distributed-Yara*. We included Distributed-Yara in our benchmark under evaluation section.

### 2.6 DREAM-Yara

DREAM-Yara is an extension of the Yara read mapper to support the DREAM framework. In DREAM-Yara, we load sequencing reads in batches and identify the bins that contain potential mapping locations for each individual reads. This is done using the IBF search distributor. As a result, we get *b* different subsets out of the loaded reads representing the different bins. A read belongs to a subset *R_bi_* if and only if it or its reverse complement or its mate pair shares enough number of k-mers with bin *B_i_*. After that, we map each subset of reads against the index of the corresponding bin. Then we collect mapping results from all subsets and consolidate them. This includes a ranking of all mapping locations per read across all bins based on mapping qualities. Finally, the mapping result is written to a single SAM/BAM file.

## 3 Evaluation

In this section, we report the results of two experiments. The first experiment focuses on evaluating the sensitivity of read mappers including that of DREAM-Yara and Yara. Our second experiment emphasizes the runtime and memory consumption of the indexing step of read mappers. We used different datasets for each experiment.

### 3.1 Index creation and updating benchmark

*Infrastructure and parameterization.* For this benchmark, all tools ran on a compute server equipped with 32 (Intel(R) Xeon(R) CPU E5-2650 v3 2.30GHz) processors and 130GB of memory. We used 8 threads whenever the tool allows parallel execution. In particular GEM, DREAM-Yara, Distributed-Yara and Bowtie-2 indices are built using 8 threads. Whereas for BWA and standard Yara a single thread is used as the indexing modules of these programs do not support parallel execution. The IBF used in DREAM-Yara is built with 18-mers and has a bit vector size of 16GB (137,438,953,472 bits).

*Datasets.* As a reasonable metagenomic reference set, we used a set of archaeal and bacterial complete genome sequences retrieved from NCBI’s RefSeq database (Haft *et al.*, 2017) dating from 2017-09-26. This dataset comprises 15,250 sequences representing 2,991 species, summing up to a total of 31.34 Gbp, a database size for which it takes significant time to compute a single index. To evaluate partial updating of the indices, the same database was selected based on the updated *Escherichia Coli* sequences from 2017-12-19. This update set sums up to 0.23 Gbp with one removed sequence and 155 new sequences, thus a typical set of sequences for which we want to update our index.

For DREAM-Yara, we partitioned the reference set into 64, 256 and 1024 independent bins using TaxSBP. As a result we get three identical reference sets that differ only in the number of bins. In the case of DIDA framework using BWA, we partitioned the reference set into 1024 parts using the provided partitioning module. Clustering times with TaxSBP are negligible, having the only cost of reading/writing the input file into separated files for each bin.

Table 1 shows time and memory required to build the index of the 31 GB reference set using different programs. It also shows how much time is needed to update the same index so that the 155 new sequences and 1 removed sequence of the species *Escherichia Coli* are accounted for. We considered Distributed-Yara, DREAM-Yara, standard Yara, Bowtie2, GEM, BWA, and DIDA framework using BWA, read mappers in this comparison. With the exception of standard Yara and BWA all the other programs were run with 8 threads. The reason is lack of support for multi-threading by the *indexing module* of the two programs. We note that direct comparison of the timing is not fair. Nevertheless, we believe that parallel building of a single big index doesn’t scale well with the number of threads. This is further supported by the results from Bowtie2 and GEM which support parallelism for indexing and were run with 8 threads.

**Table 1.** Time and memory required for building indices. Time values are wall clock times and Peak Memory refers to the maximum resident memory occupied by a program (*all threads in case of multi-threading*) during execution. Update times refer to time to rebuild indices to reflect partial changes in databases. Since there is no way to do the same for standard mappers, similar values as build time are reported[*]. The values in braces represent individual time/Peak Memory of FM-indices and IBF, respectively.

|  | bins | Build Time<br>[hr:min:s] | *Update Time<br>[hr:min:s] | Peak Memory<br>[Gb] |
| --- | --- | --- | --- | --- |
| DIDA (BWA) | 1024 | 56:07 | 2:51 | 0.06 |
| Distributed-Yara | 1024 | 49:10 | 3:15 | 8.32 |
| DREAM-Yara |  | [49:10, 17:52] | [3:15, 3:28] | [8.32, 16.15] |
|  | 1024 | 1:07:03 | 6:43 | 16.15 |
|  |  | [1:22:33, 21:28] | [6:02, 3:40] | [12.38, 16.16] |
|  | 256 | 1:44:02 | 9:42 | 16.16 |
|  |  | [1:36:56, 20:04] | [9:41, 9:23] | [16.32, 16.18] |
|  | 64 | 1:57:01 | 19:04 | 16.32 |
| Yara |  | 27:17:54 | *27:17:54 | 85.07 |
| Bowtie 2 |  | 9:51:42 | *9:51:42 | 89.80 |
| BWA-MEM |  | 19:33:24 | *19:33:24 | 43.83 |
| GEM |  | 20:41:01 | *20:41:01 | 104.00 |

DREAM-Yara requires an IBF in addition to bin-many small indices. The reported times for DREAM-Yara are the sum of time required to build/update the small indices and to create the corresponding IBF. The same is true for the reported Memory, except in this case we took the maximum instead of the sum. Distributed-Yara uses the same indices as DREAM-Yara, but there is no need for an IBF as we search all reads against all indices. So in the case of Distributed-Yara, we omit the time and memory related to IBF construction.

Distributed-Yara and the DIDA framework using BWA exhibited the best time in creating and updating indices. They also have the smallest amount memory footprint. Despite of having the least computational requirements during indexing step, we do not find them to be practical choice considering how slow they make the mapping process as it is shown in the next section. DREAM-Yara is only 18 minutes slower in creating and 3 minutes slower in updating indices than Distributed-Yara. These differences are due to the time needed for building and updating the complementing IBF.

DREAM-Yara took 1 hour and 7 minutes to index 31.34 Gb of reference sequences with 1024 bins. That is approximately 9 times faster than the next fastest indexer (Bowtie2) and 26 times faster than the slowest indexer by standard Yara. Memory consumption was lower for DREAM-Yara indexer, using 62% less memory compared against to BWA which is the next best method concerning memory. As it is shown in table 1, the peak memory consumption of DREAM-Yara indexer is coming from the construction of IBF in particular.

DREAM-Yara has also the advantage of having bin-many small indices which makes rebuilding them less time-consuming. In case of changes in reference-set, we have to rebuild only the affected indices, whereas BWA and standard Yara need to rebuild the complete index for any change in the reference set. To demonstrate the advantage of this, we considered all the reference changes under the species *E. Coli* between 2017-09-26 and 2017-12-19. This accounted for 155 new sequences and one removed sequence which affected 42, 15 and 4 bins out of 1024, 256 and 64 bins respectively. These are a small portion of the complete set and rebuilding them took 7 minutes for 1024 bins, 10 minutes for 256 bins and 19 minutes for 64 bins. This is a significant improvement considering the amount of time needed if we had one big index which is about a day for standard Yara, BWA and GEM. Whereas Bowtie2’s indexer takes about 10 hours.

### 3.2 Rabema benchmarks

*Infrastructure and parameterization.* We used the same infrastructure in this evaluation as for the indexing benchmark. For maximum throughput, all tools run using 8 threads. For DREAM-Yara and Distributed-Yara, we chose a 1024 bins scheme, as it is the fastest among 64, 256 and 1024 bins to build indices. All experiments measured read mapping throughput in *giga base pairs per hour* (Gbp/h) and memory consumption.

*Datasets.* We use a publicly released sequencing run (SRA/ENA id: SRR6504858) submitted by Nanfang Hospital of Southern

Medical University; the genomic DNA used in this study came from a human gut. This dataset consists of 2 × 150 b_p_ whole genome sequencing reads produced by an Illumina HiSeq X Ten instrument. For practical reasons, we consider only the first 40 M reads of the first pair for performance assessment and only the first 1 M reads for sensitivity analysis by Rabema.

We use the Rabema benchmark to evaluate mappers sensitivity. The *Rabema benchmark* (Holtgrewe *et al.*, 2011) (v1.2) measures the sensitivity of read mappers in finding *relevant* mapping locations of genomic reads. We parameterize Rabema to count as relevant all co-optimal mapping locations for each read, i.e., only best strata. Rabema computes the *sensitivity* of each tool as the fraction of relevant mapping locations found per read. For a thorough evaluation, Rabema classes mapping locations by their *error rate* then computes sensitivity within each error rate class. The benchmark reports two types of percentual scores, one normalized by the number of valid mappings of a read and the other absolute percentage of mapping locations without any normalization (for example, assume two reads *r*_1_ and *r*_2_ map to the database with e.g. 0 errors and assume *r*_1_ maps uniquely and *r*_2_ maps at 100 locations. Assume read mapper *m*_1_ finds all locations for *r*_1_ and *r*_2_, read mapper *m*_2_ finds the location of *r*_1_ and 50 locations for *r*_2_. In the first case Rabema will report a sensitivity of (1+1)*/*2 = 1 for *m*_1_ and (1+0.5)*/*2 = 0.75 for *m*_2_, in the second case Rabema will report for *m*_1_ a sensitivity of (1+100)*/*101 = 1 and for *m*_2_ (1 + 50)*/*101 = 0.5049.)

We built a Rabema gold standard by running RazerS 3 in full-sensitive mode within 5 % error rate. Subsequently, we provided the reads as unpaired to each tool, as the Rabema benchmark is not meaningful for paired-end reads.

Both normalized and absolute Rabema results are shown in Table 2. On the left are percentual scores normalized by the number of valid mapping locations. Hence, as pointed out above, repetitive reads have less weight. Distributed-Yara, DREAM-Yara and Yara [s=0] are the most sensitive tools in finding all co-optimal locations; They are full-sensitive all the way up to 5 % error rate. The similar sensitivity scores of all derivative mappers of Yara is an assurance for no loss in mapping sensitivity due to the distribution of mapping. GEM is not full-sensitive even though it claims to be so; it loses small fraction of normalized locations starting from 2 % error rate. Bowtie 2 and BWA are not designed for this task; indeed, they loose a number of co-optimal locations.

**Table 2.**
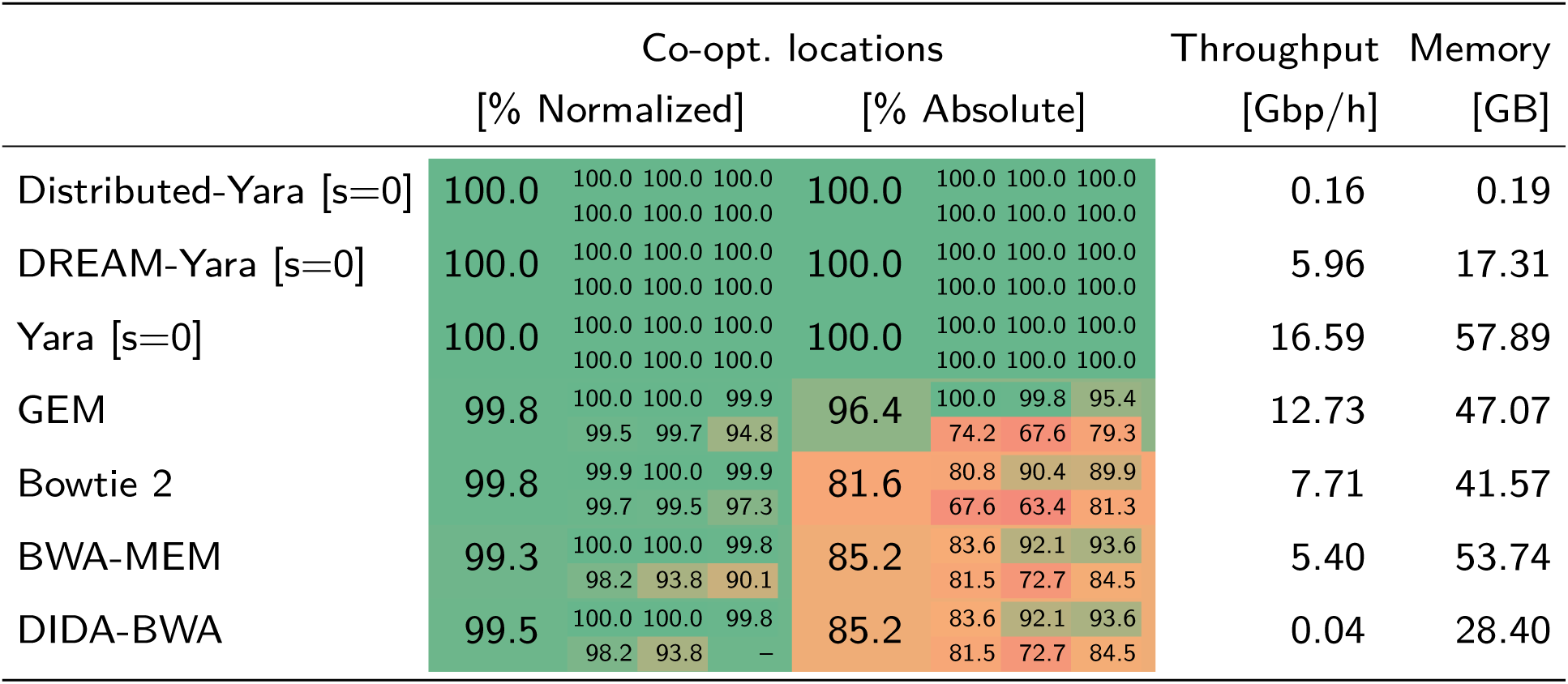
Rabema benchmark results on 1 M human gut metagenomic reads (SRA/ENA id: SRR6504858) mapped against 31.34 GB archaeal and bacterial complete genome sequences retrieved from NCBI’s RefSeq database. The colored panels show the results of finding all co-optimal mapping locations of the reads; Big numbers show total Rabema scores, while small numbers show marginal scores for the mapping locations at ^0 1 2^ % error rate. The left panel shows the sensitivity of mappers normalized by the number of locations reported per read, while the right panel shows absolute sensitivity. Memory and speed assessments on the right are done using the first 40M reads of the same read set.

However, in metagenomics “repetitive” reads often stem from multiple genomes of similar organisms as opposed to repetitive regions in a genome. Therefore all their mapping locations are significant for downstream analysis. The middle panel in Table 2 shows Rabema results where we considered the absolute number of co-optimal locations without any normalization. Here, DREAM-Yara, Distributed-Yara and Standard Yara [s=0] are the clear winners in sensitivity. They found all co-optimal locations; GEM loses 5.6 % of locations at 2 % error rate.

In addition, Yara [s=0] has the highest throughput, being 1.3 times faster than GEM, 2.15 times faster than Bowtie 2 and 3.07 times faster than BWA. The DIDA distribution framework using BWA as a core mapper is too slow and inefficient for a setup like this. It is 130 times slower than BWA with one big index. The same is true for Distributed-Yara (a naive distribution using Yara mapper.) It is 103.69 times slower than using a standard Yara mapper. DREAM-Yara, on the other hand, is only 2.78 times slower than standard Yara and competitive with the other read mappers.

In both evaluations, we showed that DREAM-Yara removes the bottleneck of large index reconstruction successfully while remaining in speed and memory consumption competitive to the standard read mappers and being 37.25 times faster than a trivial distribution. When we consider the time and space requirements for mapping, DREAM-Yara needed 56% less memory when compared against Bowtie2, BWA and standard Yara in this task.

### 4 Discussion

We hope that our work in this manuscript has an impact in various research directions. First, we think that DREAM-Yara will serve the community as a very practical, exact read mapper for Illumina reads and large databases. Second, we think that the DREAM framework will trigger further work in this area. The most crucial point in the DREAM framework is the question whether a distribution of reads can be done fast enough in comparison to the time needed for mapping the reads to one or a handful of indices. We showed that the combination of interleaved Bloom filters and the *k*-mer lemma answers this question positively.

We think it will be interesting to investigate in the future for which parameters the *k*-mer directory implementation and for which the IBF implementation have more advantages. For large, basically arbitrary *k*, we presented with the IBF a very viable way to implement an approximate search distributor.

Another interesting area to extend this work will be how to bin the sequences. How many bins are optimal for a given set? Will we benefit from allowing to further subdivide the genomic contigs? In this work we opted for a taxonomic based clustering for being a straightforward implementation. However, a clustering based on sequence similarity would be beneficial for computing the bins in DREAM-Yara. The taxonomy classification is not purely sequence-based and by definition can have distantly related groups with highly similar sequences or closely related groups with low sequence similarity. A sequence-based approach could result in a better distribution, meaning that reads would be potentially mapped to fewer bins, speeding up the mapping procedure. Important to note is that the evenness of the bin sizes makes a difference for our Bloom filter implementation and it is an important aspect to be considered when computing the binning. This comes from the fact that the IBF has bin-many Bloom filters of the *same* size.

Hence the largest bin will have the highest false positive rate for a *k*-mer, which would result in worse filtering.

Finally, it will be interesting to investigate whether other indices can take advantage of the framework. While FM-indices support fast approximate searching, they do not support the compression of similar sequences. However, the bins in the DREAM index will contain by construction very similar sequences. Hence we anticipate that compressed indices together with the IBF based search distributor will result in very practical, space efficient and fast read mappers for very large, repetitive databases.

### 5 Conclusion

In conclusion, we presented the novel DREAM framework for distributed read mapping which can support fast updates of the sub-indices as well as compression by design. We implemented within this framework DREAM-Yara, a distributed version of the Yara (Siragusa, 2013) read mapper. The main contribution lies in our implementation of the dynamic search distributor by using interleaved Bloom filters together with the *k*-mer counting lemma and an in-memory distributed version of Yara. We showed that the resulting read mapper is very competitive in terms of speed, it is faster than BWA and only slightly slower than the Bowtie2, which both use one large FM-index, but more than 37.25 times faster than a trivial distribution. We also showed that DREAM-Yara can conduct a typical batch index update for a metagenomic dataset in about 6 minutes, whereas the rebuilding Bowtie2 index takes 10 hours and that of BWA or Yara index takes about a day.

DREAM-Yara is part of the SeqAn library for efficient data types and algorithms.

## Acknowledgments

We acknowledge Mikaël Salson for his ideas in defining the DREAM framework jointly with Knut Reinert.

## Funding

This work was supported by the Coordenação de Aperfeiçoamento de Pessoal de Nível Superior (CAPES) - Ciência sem Fronteiras (BEX 13472/13-5) to VCP, the InfectControl 2020 Project (TFP-TV4), and the BMG Project "Metagenome Analysis Tool" (2515NIK043). The authors also acknowledge the support of the de.NBI network for bioinformatics infrastructure, the Intel SeqAn IPCC and the IMPRS for Scientific Computing and Computational Biology.

